# Acute oral toxicity of zinc phosphide: an assessment for wild house mice in Australia

**DOI:** 10.1101/2021.12.09.472031

**Authors:** Lyn A. Hinds, Steve Henry, Nikki Van de Weyer, Freya Robinson, Wendy A. Ruscoe, Peter R. Brown

**Affiliations:** CSIRO Health and Biosecurity, GPO Box 1700, Canberra, Australia, 2601

**Keywords:** *Mus musculus domesticus*, acute rodenticide, LD_50_, LD_90_, outbred laboratory mice, efficacy, aversion

## Abstract

**BACKGROUND:** The efficacy of zinc phosphide (ZnP) for broadacre control of wild house mice in Australia is being reported by growers as increasingly variable. Have mice become less sensitive over time or are they taking a sub-lethal dose and developing aversion? In this laboratory study the sensitivity of groups of wild caught and an outbred laboratory strain of mice was assessed using oral gavage of a range of ZnP concentrations. The willingness of mice to consume ZnP-coated grains was then determined.

**RESULTS:** Each mouse group had very similar LD50 values (72 to 79 mg ZnP per kg body weight) which are significantly higher than previously reported. Time to death post-gavage ranged between 2.5 to 48 h. ZnP coated grains (50 g ZnP per kg grain) presented in the absence of alternative food were consumed and 94 percent of wild mice died. Mice provided with alternative food and ZnP coated wheat grains (either 25 or 50 g ZnP per kg grain) consumed toxic and non-toxic grains, and mortality was lower (33 to 55 percent). If a sublethal amount of ZnP coated grain was consumed, aversion occurred mostly in the presence of alternative food.

**CONCLUSIONS:** The sensitivity of wild house mice to ZnP in Australia is significantly lower than previously assumed. Under laboratory conditions ZnP coated grains coated with a new higher dose (50 g ZnP per kg grain) were readily consumed. Consumption of toxic grain occurred when alternative food was available but was decreased. It is important to assess the efficacy of the higher ZnP dose per grain under natural field conditions, especially when background food is low.

## 1 INTRODUCTION

Zinc phosphide (Zn_3_P_2_, hereafter ZnP) has been used for the control of several commensal and field rodents for several decades.^1^ It has the advantage of being an acute rodenticide with low risk of secondary poisoning and an absence of long-term residues,^2, 3^, but it is not rodent specific. Many other mammals (ungulates, carnivores, lagomorphs) and birds (particularly land and waterfowl and granivorous passerines) are sensitive to ZnP and fish are highly susceptible.^4–6^ The mode of action of ZnP is primarily due to the release of phosphine gas and its subsequent uptake into the blood stream and major organs. When food enters the stomach it generally triggers an increase in the release of gastric acid.^7, 8^ After consumption of ZnP-food baits the hydrolysis of ZnP results in a rapid release of phosphine which causes acute toxicity in various organs including the lungs, liver and heart. Animals show adverse changes in respiratory patterns, hypoxia and loss of control of bodily movements (ataxia). In the absence of food in the stomach undegraded ZnP can be absorbed systemically, and animals may survive beyond 48 h but succumb to liver and/or renal failure 5 - 14 days later. ZnP is thought to disrupt cytochrome C oxidase, leading to formation of highly reactive oxygen compounds, which result in tissue injury, particularly those with a high oxygen demand (brain, lungs, liver, and kidney).^8^

The ZnP oral toxicity (LD_50_ - lethal dose that kills 50% animals) for mammals ranges from 5.6 mg/kg for nutria (*Myocaster coypus*)^4^ and 9.6 mg/kg for brushtail possums (*Trichosurus vulpecula*)^9^ to 60-70 mg/kg for sheep, *Ovis aries.*^5^ For several rodent species the LD_50_ is in the range of 20-40 mg ZnP/kg body weight.^4, 6^ Specifically, for wild house mice, *Mus musculus*, the LD_50_ is reported as 32.7 mg/kg body weight, whilst for Swiss Webster albino mice, an outbred strain of laboratory mice, the value is 54 mg/kg body weight,^10^ though Bell^11^ reported a value of 25 mg/kg for white mice.

The wild house mouse (*Mus musculus domesticus*) is a major pest in grain production areas of Australia.^12, 13^ Mouse outbreaks occur irregularly and cause significant economic losses and social impacts.^12–18^ ZnP was first registered in 2000 as an in-crop acute rodenticide for mouse management^19, 20^ and initially was applied to grain crops at 1 kg/ha (registered rate of 25 g ZnP/kg wheat; ∼3 toxic grains/m^2^; each grain is coated with approximately 1 mg - the equivalent of an LD_90_ dose for wild mice) during periods of increasing or high mouse numbers. Its efficacy was highest when applied when food was scarce, especially prior to crop/cereal flowering.^13, 18^ After aerial application there was a 51% reduction in adjusted trap success for mice, while after perimeter baiting less than 25% reduction was observed.^21^ In some situations where outbreaks are frequent, it has become general practice for farmers to apply ZnP as a preventative control strategy when a crop is being sown, even if mouse numbers are low and not predicted to increase (Henry S and Brown PR unpublished observations). Furthermore, several applications of ZnP bait have been required during the crop growing season leading some farmers to question whether there has been a change in the effectiveness of ZnP over the last 10-15 years (Henry S and Brown PR unpublished observations). This may also be influenced by the broadscale adoption of conservation agriculture practices which provide increased cover and reduced soil disturbance.^22^ These observations have raised questions such as: (1) have mice become tolerant to the toxin? (2) has presentation of the toxin on cereal grain led to higher levels of bait aversion? and (3) does the presence of moderate background food resources reduce uptake of ZnP-coated grains which are spread at approximately 3 grains/m^2^?

The effectiveness of ZnP-coated grains has previously been assessed in the laboratory in the presence of alternative food.^23^ In that study, wild mice showed a strong preference for consumption of cereal grains over lentils, and this influenced their uptake of toxic ZnP-coated barley grains. The mortality rate for mice presented with a choice of lentils and toxic ZnP-coated cereal grains was 86%, whereas for mice presented with a choice of cereal grains and toxic ZnP-coated cereal grains the mortality rate was only 50%. One possible interpretation here is that the presence or absence of food in the stomach of mice directly impacts the efficacy of the ZnP. Another possible change in the two decades since ZnP has been registered for control of mice in crops in Australia is that regular and repeated bait applications may have led to the selection of mice that are now less sensitive or more tolerant to ZnP. Potentially, mice in areas where ZnP has been applied on a regular basis may be more tolerant to ZnP than mice that have never been exposed to the toxin.

The current laboratory study was designed to re-examine the acute toxicity of ZnP in fed and unfed house mice. Our first objective was to determine the LD_50_ for the wild house mouse in Australia using oral gavage of known concentrations of ZnP in vegetable oil. To achieve this, wild mice were trapped from an area where ZnP had been used most years, and from a location where ZnP had never been used. Their responses were compared with those of an outbred laboratory strain of mice. Considering the results obtained we then assessed whether wild caught house mice would consume wheat grains coated with a higher dose of ZnP than is currently in use.

## 2 METHODS

### 2.1 Animals

Wild house mice were trapped using single live capture traps (Longworth, Abingdon, UK). Traps were set with wheat grains and non-absorbent Dacron in the late afternoon and checked from dawn the following morning. Trapped mice were weighed, measured, sexed and assessed for general condition before being transferred to small cages (30.5 x 13.5 x 12.0 cm, length x width x depth) containing wood shavings, nesting material, a cardboard tube, apple pieces for moisture, a mounted water bottle and food pellets (standard maintenance diet rat and mouse pellets) (Gordon’s Specialty Stockfeeds, NSW, Australia). Mice were transported to the animal facility at CSIRO, ACT, Australia, in an air-conditioned vehicle. On arrival, they were transferred to individual, clear, mouse-specific cages (26 x 40 x 20 cm, length x width x depth) which contained bedding and nesting material (wood shavings and tissue paper), water and food *ad libitum*, cardboard tube shelters and popsicle sticks for environmental enrichment. The room temperature was maintained at 22 ± 2° C. During the two-week acclimation period mice were checked daily for alertness, and hydration, and weighed twice per week.

### 2.2 Experimental design

#### Experiment 1: Determination of the LD_50_ and LD_90_ of wild and laboratory house mice

Wild house mice were trapped from a farm location on Yorke Peninsula, South Australia, where ZnP baiting occurred regularly (at least once per year for 9 of the last 12 years). This group is hereafter referred to as wild mice (Exposed). The second group of wild house mice was trapped from around a horse stable in Murrumbateman, NSW, where ZnP baiting has never occurred. This group is hereafter referred to as wild mice (Naïve). The third group of mice, an outbred strain of laboratory mice, was obtained from the Animal Resources Centre, Western Australia. Within each experimental group equal numbers of males and females were used when available.

We used oral gavage of a range of concentrations of ZnP in vegetable oil using the broad principles of the OECD Up Down protocol (UDP).^24, 25^ Starting doses were based on the reported LD_50_ values.^10^ For the outbred laboratory strain the starting dose was 50.22 mg ZnP/kg body weight, for the wild caught mice it was 30 mg ZnP/kg body weight. If any dosed mice survived, a higher dose was administered to new individuals, while if they died a lower dose was administered to new individuals. This process was repeated until a dose response curve was established. Doses given ranged between 30 - 149 mg ZnP/g body weight.

Specific doses based on body weight were given to animals lightly anaesthetised with Isoflurane (4%) in oxygen, using a 20 g curved gavage needle attached to a 1 ml syringe between 08:00 and 09:00 h each morning. Once the gavage needle was in position the plunger of the syringe was gently depressed to administer the individual’s calculated dose (90-300 µl) into the stomach. The gavage needle was gently removed, and the animal maintained in a vertical position for a few seconds until it showed the swallow reflex and signs of recovery from the anaesthetic. Each mouse was placed in a glass observation tank (30 x 30 x 30 cm) covered with a wire mesh lid. The oral gavage procedure took less than 20-30 seconds and anaesthetic recovery was rapid and uneventful.

Animals were observed regularly (every 30 min) for 24 h post gavage and more frequently (every 15 min) if signs of toxicity were observed. Observation of their behaviour included: general alertness (normal to lethargic) and movement (normal exploratory activity to abnormal/unbalanced and a reluctance to move); body position (normal to sternal or lateral recumbency); clarity of eyes (bright, clear and open to dull, half or fully closed); respiratory pattern (normal, shallow; gasping; slow, rapid). If mice exhibited lateral or sternal recumbency and had lost their righting reflex, as well as showing high respiration rate and lethargy, they were humanely killed in accordance with animal ethics requirements for this project.

Mice allocated to a treatment dose were divided into a fed or unfed group. For the fed group, food was available *ad libitum*, while for the unfed group, food was removed 12 h prior to oral gavage with a ZnP dose. Food was returned to unfed individuals at 8 h post gavage. Dosed mice were checked through to 48 h post gavage as per the UDP^25^ and then humanely killed using isoflurane then cervical dislocation.

#### Experiment 2: Determination of uptake and efficacy of ZnP-coated grain (LD_90_ dose defined in Experiment 1)

Wild caught house mice were trapped from around Parkes, NSW, and transported back to the laboratory. They were acclimatised for 7-10 days in individual cages and provided with a maintenance diet of laboratory mouse pellets and wheat grains prior to the experiment, as described above. Three days before the start of the experiment, laboratory chow was removed, and only standard wheat grains (hereafter referred to as alternative food) were offered.

We established three experimental groups. Firstly, we compared intake and mortality of one group of animals provided with a choice of alternative food (“Fed”) and ZnP-coated grains (50 g ZnP/kg wheat grain) (hereafter referred to as F50), with another group in which only ZnP-coated grains (50 g ZnP/kg grain) (“UnFed”; hereafter UF50) were provided for 4 h before the addition of alternative food.

Secondly, a third group of animals was provided with a choice of alternative food and ZnP-coated grains (25 g ZnP/kg grain) (hereafter F25). We did this to confirm results obtained for animals receiving a choice of ZnP-coated grains (25 g ZnP/kg grain) and alternative food as described in ^23^ and to compare this F25 group with the above-mentioned F50 group. Two sequential independent trials (Trial 1 and Trial 2), each using ten animals per treatment group, were undertaken over a two-week period. The preparation of batches of ZnP-coated wheat is described below.

Individual mice were presented with two small dishes, one containing wheat grains (≥15% body weight, (g)) and one containing ZnP-coated wheat grains (*n* = 10). This choice was offered for three consecutive days. Each day, consumption of each food type was recorded, and the dish positions reversed in individual mouse boxes. Each dish was secured to the floor of the cage with Blu Tack (Bostick Pty Ltd, Thomastown, Victoria, Australia) to minimise spillage.

Mice were monitored at 30 min intervals from the time of introduction of the ZnP-coated wheat grains for 24 h (Days 1 - 3) and then 2-3 times per day between Days 4 - 6. Mice that were still alive on Day 6 were humanely killed as described above.

On Days 1 - 3, between 08:00 - 09:00 h, food was removed, cages were cleaned, and animals were weighed and checked for any signs of distress and returned to the cage. Then at 16:00 h, for the fed groups (F50 and F25), alternative food (≥15% body weight) and their respective ZnP-coated grains (*n* = 10) were provided in separate dishes and observations commenced. At this time, 16:00 h, the unfed group (UF50) was provided with ZnP-coated grains (*n* = 10) only and observations commenced. Their alternative food (≥15% body weight) was added between 20:00 and 22:00 h.

To potentially increase uptake of ZnP-coated grains, in the second trial no alternative food was provided to any groups until approximately 6 h after provisioning of ZnP-coated grains on Days 2 and 3.

### 2.3 Zinc Phosphide

#### 2.3.1 Preparation of concentrations of ZnP in oil and coating of ZnP onto grains

For Experiment 1, the ZnP powder (93% purity; ≥ 19% active phosphor (P) basis, powder; Lot # STBJ1895; Sigma Aldrich Castle Hill NSW, Australia) was weighed and added to 100% pure vegetable oil (Crisco, Goodman Fielder, Moorebank, NSW, Australia) to achieve a known concentration (range: 3.22 - 20 mg/ml) for oral gavage on that day. The powdered ZnP in oil forms a suspension of ZnP particles and is not a homogenous solution. Therefore, the prepared suspension was rapidly vortexed before each individual dose was quickly loaded into the 1 ml syringe for gavaging. Each day, two aliquots of each concentration were removed, held at 4° C, and subsequently submitted for independent analysis (see below).

For Experiment 2, two batches of ZnP-coated wheat grains were prepared in-house. following the method used by commercial bait producers who pre-mix ZnP powder and sterilised grain before adding oil. To achieve the rate of 25 g ZnP/kg grain, 250 g wheat was weighed, and 7.8 g ZnP powder (corrected for 80% purity) was added to a glass bottle. The bottle was capped and continuously rotated until all the powder was distributed on the grains after which 6.7 ml vegetable oil was added. The rotating process was repeated until all the grains were evenly covered with both powder and oil.

To achieve the rate of 50 g ZnP/kg grain, the above procedure was repeated using 125 g wheat grains, 7.8 g ZnP powder and 3.0 ml vegetable oil.

Coated grains were then spread onto trays on aluminium foil and placed in a drying oven at 40° C overnight. ZnP-coated grains were then returned to a labelled airtight glass bottle and stored in a dark cupboard.

ZnP-coated grains (n= 8-10) from each batch were individually analysed to independently confirm the expected coating rate of approximately 1 and 2 mg ZnP per grain, respectively.

#### 2.3.2 Analysis of concentrations of ZnP in oil and ZnP on coated grains

Analysis was undertaken by ACS Laboratories (Australia) (37 Stubbs St, Kensington, Victoria, 3031), a commercial, independent analytical service provider accredited by the National Association of Testing Authorities (NATA), Australia (NATA Accreditation No 16973). One aliquot of each ZnP concentration in oil was subsampled (in duplicate) and analysed by microwave digestion followed by inductively coupled plasma optical emission spectroscopy (ICP-OES). Approximately 200 μl subsamples were weighed directly into Teflon digestion tubes to which 5 ml of concentrated nitric acid and 1 ml of 30% hydrogen peroxide were added. Grains coated with ZnP were also individually analysed by microwave digestion followed by ICP-OES. Each grain was weighed directly into a Teflon digestion tube to which 5 ml of concentrated nitric acid and 1 ml of 30% hydrogen peroxide were added. Each digestion tube was then microwaved for 65 minutes using a temperature gradient from 0-200° C in a Milestone Connect Ethos Lean Compact Microwave Digestor. Once cooled the contents of the digestion tube were diluted to 50 ml in deionised water, followed by a further 10-fold dilution in deionised water for ICP analysis using an Agilent 5110 ICP-OES (Dual View). The operating conditions for the instrument were: 1.20 kW RF power; plasma 15.0L/min; auxillary 1.50 L/min; AVS7 sample injection system; injection pump rate 15.7 ml/min; detection - intensity, Zn 206.2 nm/c/s.

The recovery of Zinc measured across a range of known concentrations of ZnP in oil (7.3, 15.5, 21.4 mg/ml) (quality controls prepared by ACS laboratories) was 100.4 ± 5.4% (n=4), 95.4 ± 7.9% (n=3) and 101.3 ± 3.5% (n=6) (Mean ± SEM) respectively. The concentrations of ZnP in oil used for oral gavage were confirmed to be within 96.3 ± 3.7% (Mean ± SEM) (n=16 samples) of that expected. The independent analysis of the two batches of ZnP-coated grains confirmed the expected average coating rate per grain. On average individual grains from the batch mixed at 25 g ZnP/kg grain were coated with 1.06 ± 0.11 mg ZnP (Mean ± SEM; n=8) (1.7, 1.0, 0.9, 1.3, 0.7, 0.9, 1.0, 1.0 mg ZnP/grain), those from the batch mixed at 50 g ZnP/kg grain were coated with 2.4 ± 0.37 mg ZnP (Mean ± SEM; n=10) (1.6, 2.9, 2.2, 1.9, 4.8, 3.6, 1.3, 2.0, 1.3 mg ZnP/grain).

## 2.4 Statistical analysis

The results of the oral gavage were assessed using generalized linear models (GLM), performed in R.^26^ A GLM, was performed using the ‘*lme4’* package^27^ specifying a binominal error structure to investigate differences in the proportion of mice that died within each mouse group and dose. To investigate the additional effects that group, sex, fed or unfed, and their interactions had on the proportion (p) of animals that died, model Akaike information criterion (AIC) values were compared between the fitted models. The estimated LD_50_ (where p = 0.5) and LD_90_ (p = 0.9) values and standard error (SE) for each group were calculated from the best fit model using the *dose.p* function in the *MASS* package.^28^ Estimated LD_50_ and LD_90_ were completed for each mouse group separately and plotted using the ‘*plot*’ function. Means are presented ± 1 standard error (SEM) throughout.

## 3 RESULTS

### 3.1 Experiment 1: Determination of the LD_50_ and LD_90_ of wild and laboratory house mice

The onset of clinical signs of toxicity varied from 2 h to > 6 h post gavage. The time to death (humane killing) in mice ranged from 2.5 h to 48 h (Table 1). Animals initially showed no signs of toxicity for several hours but then succumbed within 30-60 min. The first sign of developing toxicity was a change from a normal resting posture to a hunched posture, increasing prominence of the guard hairs (pilo-erection) and rapid, shallow breathing. This was followed by an increased reluctance to move when disturbed, and/or moving with an unsteady gait (ataxia), and the eyes were half-closed. Animals then showed lateral or sternal recumbency and loss of righting reflex at which point they were humanely killed. In general, the higher the oral gavage dose the more rapid the onset of signs of toxicity and the shorter time to death (Table 1) (Table S1 shows individual timing).

**Table 1:** Mortality (%) and time to death (h) for different mouse groups after oral gavage of different concentrations of ZnP in oil.

| Dose<br>(mg ZnP/kg) | *Body weight<br>(g) | n | Mortality<br>(%) | Time to death<br>(h post gavage) (range) |
| --- | --- | --- | --- | --- |
| **Outbred laboratory mice** |  |  |  |  |
| 50.22 | 29.9 ± 1.8 | 8 | 25.0 | 25.0 - 25.3 |
| 65.1 | 34.9 ± 1.6 | 6 | 33.3 | 31.0 - 48.0 |
| 74.4 | 34.8 ± 1.5 | 5 | 0.0 | - |
| 83.7 | 32.2 ± 1.8 | 8 | 62.5 | 9.3 - 29.5 |
| 102.3 | 31.9 ± 1.8 | 8 | 87.5 | 7.5 - 28.0 |
| 148.8 | 32.0 ± 2.1 | 4 | 100.0 | 6.7 - 10.0 |
| Mean ± SEM | 32.4 ± 0.7 (n = 39) |  |  | 18.5 ± 2.4 (n = 20) |
| **Wild mice (Naïve)** |  |  |  |  |
| 29.95 | 12.9 ± 1.8 | 8 | 0.0 | - |
| 50.22 | 15.3 ± 1.8 | 8 | 37.5 | 6.75 - 23.8 |
| 74.4 | 15.3 ± 1.2 | 7 | 71.4 | 6.3 - 19.0 |
| 102.3 | 13.3 ± 1.8 | 8 | 62.5 | 3.17 - 8.4 |
| 148.8 | 15.4 ± 1.8 | 8 | 100.0 | 2.5 - 24.5 |
| Mean ± SEM | 14.5 ± 0.7 (n = 39) |  |  | 9.8 ± 1.5 (n = 21) |
| **Wild mice (exposed)** |  |  |  |  |
| 30.41 | 16.3 ± 0.8 | 8 | 0.0 |  |
| 50.22 | 10.9 ± 0.7 | 7 | 14.3 | 24.0 |
| 65.1 | 11.1 ± 0.9 | 4 | 25.0 | 19.8 |
| 74.4 | 11.6 ± 0.8 | 8 | 62.5 | 3.5 - 23.3 |
| 83.7 | 9.9 ± 0.8 | 6 | 50.0 | 6.7 - 32.7 |
| 102.3 | 12.4 ± 0.8 | 11 | 90.9 | 4.0 - 24.0 |
| Mean ± SEM | 12.3 ± 0.5(n = 44) |  |  | 11.8 ± 2.0 (n = 20) |

\*Body weight (Mean ± SEM) of dosed animals on day of treatment, includes males and females.

For the laboratory outbred mice (average body weight = 32.4 ± 0.7 g), the time to first onset of signs of toxicity was delayed compared to the two groups of wild house mice (average body weight for naïve and exposed mice = 14.5 ± 0.7 and 12.3 ± 0.5 g respectively). The time to death post gavage was also variable both within and between mouse groups, ranging from 6.7 - 48 h in outbred lab mice, 2.5 - 24.5 h in naïve wild mice, and to 3.5 - 32.7 h in exposed wild mice (Table 1).

The proportion of mice that died was dose-dependent (Fig. 1). The proportion of animals that died at each treatment dose was not different between lab and wild groups, sex, or fed and unfed status, therefore adding these factors did not improve the model fit (ΔAIC= 6.43; Table S2).

**Figure 1:**
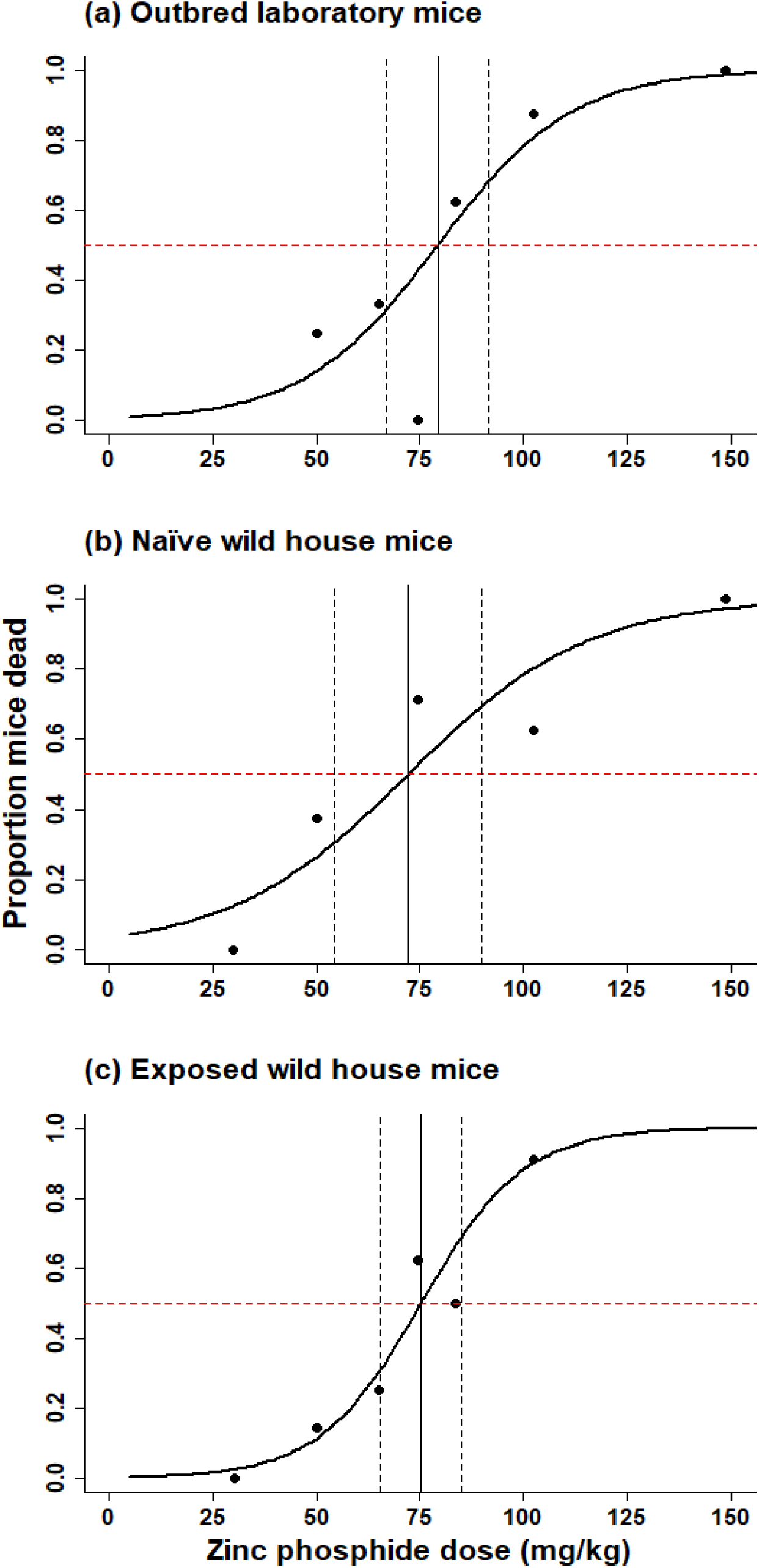
Proportion of mice dying after oral gavage with different ZnP concentrations (mg ZnP/kg body weight). Calculated dose response curves for (a) Outbred laboratory mice (b) Naïve wild house mice and (c) Exposed wild house mice. Horizontal red dashed line represents 50% mortality, vertical solid line equates to LD_50_ value, vertical dashed lines represent standard error for the LD_50_ estimate. n>4 animals per test dose, with a mix of males and females.

**Table 2:** Estimated LD_50_ and LD_90_ values (mg ZnP/kg body weight; Mean ± SEM) for different groups of mice.

| Mouse group | Total<br>number<br>mice<br>dosed | LD<sub>50</sub><br>(mg ZnP/kg body weight) | LD<sub>90</sub><br>(mg ZnP/kg body weight) |
| --- | --- | --- | --- |
| Outbred Laboratory strain | 39 | 79.18 ± 6.24 | 114.58 ± 14.6 |
| Naïve wild | 39 | 72.11 ± 9.09 | 119.50 ± 18.01 |
| Exposed wild | 44 | 75.22 ± 4.39 | 102.27 ± 9.34 |

The estimated LD_50_ values for each mouse group ranged between 72 and 79 mg ZnP/kg body weight (Fig. 1; Table 2; P>0.05). The estimated LD_90_ values also were not significantly different between the different mouse groups, ranging between 102 and 120 mg ZnP/kg body weight; Table 2; P>0.05).

None of the Outbred Laboratory mice dosed at 74.4 mg ZnP/kg body weight died (Fig. 1a; Table S1). If this treatment dose was not included in the model estimating the LD_50_ and LD_90_ values for outbred mice, the new estimates were 72.8 ± 7.1 and 109.6 ± 14.6 mg ZnP/kg body weight respectively which are not significantly different from the original estimates (Table 2).

### 3.2 Experiment 2: Uptake of ZnP-coated wheat grains

Across both Trials and all treatment groups, mice consumed ZnP-coated grains (18 of 20 mice in UF50, 15 of 20 mice in F50, and 19 of 20 mice in F25) (Table 3), predominantly on the first day of presentation (Tables S3 and S4). Three of the 60 mice (one in each of F50 T1, UF50 T1 and UF50 T2) never consumed any ZnP-coated grains on any of the three days and have been excluded from the analyses.

**Table 3:**
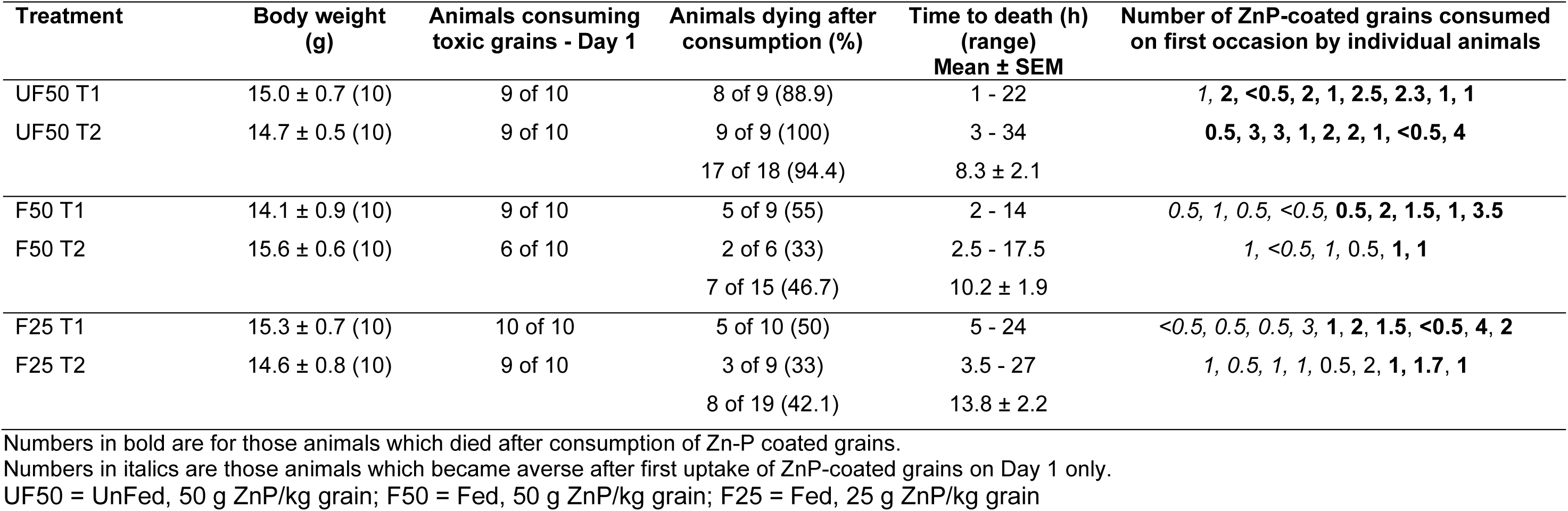
Responses of wild mice on Day 1 to exposure and uptake of ZnP-coated grains for each treatment over two trials (T1, T2). Body weights (Mean ± SEM (n)) of each group at start of each treatment round, number of animals consuming ZnP-coated grains on Day 1, mortality (%) after consumption, time to death (range, Mean ± SEM), and number of ZnP-coated grains consumed by individual mice.

For Treatment UF50, nine of 18 animals (50%) were observed to consume ZnP-coated grains within 30-60 mins of their presentation. For the remaining nine animals, six consumed ZnP-coated grains before alternative food was added, while three animals consumed ZnP-coated grains between 1 and 3 h after the addition of alternative food.

For the F50 and F25 treatments in which there was simultaneous presentation of the ZnP-coated grains with alternative food, two of 15 F50 animals (13.3%) and six of 19 F25 animals (31.5%) consumed ZnP-coated grains within the first 60 min. The remaining animals in these fed groups (86.5% and 68.5% respectively) consumed ZnP-coated grains between 2.5 and 19 h after presentation.

Across all treatments, after first recorded consumption of ZnP-coated grain, the shortest time to the onset of signs of toxicity and subsequent humane intervention was 30 min and 1 h respectively. The longest interval to intervention after first recorded ZnP-coated grain consumption was 34 h. The average time to intervention for the three treatments was similar: 8.3 h for UF50,10.2 h for F50 and 13.8 h for F25 (Table 3).

For the two trials of UF50 treatments in which alternative food was not available for the first 4-6 h after presentation of ZnP-coated grains, 17 of 18 mice that consumed ZnP-coated grains died (∼94%). By comparison, for the two trials of the F50 treatment where alternative food was available, 15 of 20 mice consumed ZnP-coated grains on Day 1 and seven of those died (∼47%) (Table 3). Similarly, over the two trials of the F25 treatment, on Day 1, eight of 19 mice that consumed ZnP-coated grains died (42%) (Table 3). Irrespective of treatment, the number of ZnP-coated grains consumed varied from <0.5 to 4, usually 1-2 grains (Table 3).

For Trial 2, on Days 2 and 3, the F50 and F25 groups did not receive alternative food for 6 h after presentation of ZnP-coated grains (i.e. same unfed treatment as the UF50 group). For F25 on Day 2, three of the seven remaining mice consumed ZnP-coated grains (*n* = 5 - 8 coated grains) before the addition of alternative food and died, but no further consumption or deaths occurred on Day 3 (*n* = 4 mice). For F50 on Day 2, four of the eight remaining mice consumed grains (0.5 - 2 ZnP-coated grains) and died, and on Day 3, one of the remaining four mice consumed 2.5 ZnP-coated grains and died (Table S4).

**Table 4.** Mice showing aversion after first consumption of Zn-P coated grains across Trials 1 and 2

| Treatment |  | n | Animals (n) that did not die after taking ZnP-coated grains on Day 1 and did not consume thereafter | Averse (%) |
| --- | --- | --- | --- | --- |
| UF50 | T1, T2 | 9, 9 | 1, 0 | 6 |
| F50 | T1, T2 | 9, 6 | 4, 3 | 46.7 |
| F25 | T1, T2 | 10, 9 | 4, 4 | 42.1 |
| Totals |  | 52 | 16 |  |

Some animals (*n* = 16/52) which consumed all or part of a ZnP-coated grain on Day 1 (Table 4), did not show any signs of toxicity, did not die, and did not consume ZnP-coated grains on either Days 2 or 3. This aversion was higher in the groups where background wheat grains were presented simultaneously with the ZnP-coated grains (F25, n=8, aversion = 42%; F50, n=7, aversion = 46.7%; UF50, n=1, aversion = 6%) which reduced the quantity of toxic grain taken (Table 4).

## 4 DISCUSSION

This assessment of the acute oral toxicity of ZnP has clearly demonstrated that wild house mice in Australia are significantly less sensitive (LD_50_ of 72-75 mg ZnP/kg body weight) than previously reported for wild mice in the USA (LD_50_ of 32.68 mg ZnP/kg body weight)._10,11_ The latter value was used by the Australian Pesticides and Veterinary Medicines Authority (APVMA) to define the dose rate (25 g ZnP/kg grain, whereby ∼1 mg ZnP per grain delivers an LD_90_ to a 15 g mouse) for registration of ZnP for broadscale management of house mice in cereal crops,^19, 20^ Our findings indicate that a higher loading of ∼2 mg ZnP/grain (equivalent to an LD_90_ loading) would be required to achieve a lethal outcome after the first consumption of a single ZnP-coated grain.

In our oral gavage study the outbred strain of laboratory mice also showed a similar LD_50_ of 79.18 mg/kg to that of the wild mice, whereas Li and Marsh^10^, using Swiss Webster outbred mice, reported an average LD_50_ value of 53.34 mg/kg. They also observed a significant difference between genders reporting values for males and females of 73.23 mg/kg and 38.86 mg/kg body weight respectively. We detected no gender differences in LD_50_ estimates within any mouse group, and the LD_50_ values were all within the same range for the three different mouse groups (72-79 mg/kg). Given the similarity in the LD_50_ values for naïve and exposed wild mice this also suggests that in the last 20 years regular field application of ZnP has not led to selection either for non-responders or for tolerant individuals. There have been no previous reports in the literature indicating any development of tolerance or genetic resistance to ZnP.^29^

Nevertheless, we observed high variability in the acute response for both outbred laboratory mice and wild mice after oral gavage intake of ZnP in each of our experiments. For example, at the estimated LD_50_ of ∼74 mg ZnP/kg body weight, 60-70% exposed and naïve wild mice died whereas none of the outbred laboratory mice died at this dose. In their original study,^10^ 80% wild mice dosed at 35.62 mg ZnP/kg body weight died whereas none of our wild mice (either exposed or naïve) died after dosing at 30 mg ZnP/kg body weight.

The earliest time at which our study animals were humanely killed after their oral gavage dose of ZnP or after consumption of ZnP-coated grains was one hour but could be up to 48h. This time-to-death range and the development of various signs of acute toxicity are similar to that observed in other studies in mice and rats.^23, 30^ This reflects the expected effects of phosphine gas production in the gut and/or direct uptake of ZnP across the gut and subsequent failure of major internal organs such as the liver, kidney, brain and heart.^8^

When wild mice were presented with ZnP-coated wheat grains (either ZnP50 or ZnP25), there was also considerable variation in responses after consumption including a rapid development of aversion as previously reported for several species including wild mice^23^, common voles, *Microtus arvalis*^31^ and prairie voles (*Microtus ochragaster* Wagner).^32^ Any wild mouse taking a sub-lethal dose on first exposure, rarely consumed ZnP-coated grains subsequently and this aversive response was higher when there was a choice of alternative food (42 - 47% mice became averse after consuming ZnP25- or ZnP50-coated grains). Aversion was very low (6%) when ZnP50-coated grains were the only food choice present for >4 hours but this is because 94% mice died after their first consumption of ZnP50 grains. We therefore conclude that increasing the ZnP dose per grain does not decrease its palatability with aversion likely due to the sub lethal adverse effects of the toxin which is much more likely to occur with the ZnP25-coated grains.

*In vitro* digestion studies have shown that the production of phosphine gas was reduced where wheat grain was digested prior to the addition of ZnP-coated grains.^33^ It was concluded that this reduction was most likely due to the inhibition of degradation of the ZnP in the presence of digested wheat grains and gastric enzymes. If this occurred in mice, the presence of food in the stomach could affect the outcome of ZnP-coated grain consumption. However, we have no direct evidence of this occurring in either of our experiments. Our observations suggest the variation in responses is due to a combination of different factors rather than a single factor - we suggest that the outcome for individual mice might vary due to quantity of ZnP consumed at first presentation (the number of grains consumed and the loading per grain), the presence of alternative food, their physiological response in the gut and their body weight.^30^

The amount of ZnP coating individual wheat grains was independently analysed and, on average, matched the expected dose rate (either 25 g ZnP/kg grain or 50 g ZnP/kg grain). However, there was variation between grains in their ZnP loading which could have affected mortality rates and aversion particularly when consumption of these ZnP coated grains occurred in the presence of other food. Similar variations were observed for a commercially prepared batch of grains used in another study^23^ and may be due to varying amounts of ZnP powder being trapped in the crease of grains of different sizes and shapes during coating.

In Australia poisoning of birds after ZnP baiting has rarely been recorded ^13, 18, 21^. If there is a higher ZnP coating per grain, does this represent a greater risk to non-target species, such as granivorous birds? For the currently registered rate, granivorous birds would need to detect and consume many ZnP-coated grains before a lethal dose accumulated (see Table 26^29^). Whether doubling the dose of ZnP coating per grain alters the potential for non-target impacts in field situations is not known and must be carefully examined. However, given that ZnP-coated grains can only be applied in-crop in Australia non-target poisoning of wild birds should remain infrequent.^21, 33^ It is also unlikely that non-target species would be at greater risk of secondary poisoning as ZnP does not accumulate in tissues and the greatest risk would be if the predator consumed the digestive tract which may contain undigested ZnP.^2^

As noted earlier, given the increased LD_50_ value for ZnP in wild mice, the coating per grain which represents a lethal dose (LD_90_) for most 15 g mice needs to be approximately 2 mg ZnP per grain. This increased coating rate means that the majority of ZnP-coated grains presented to mice would likely be a lethal dose, effectively reducing the chances of mice encountering a sub lethal dose and developing aversion. When wild mice were presented with grains coated with ∼2 mg ZnP they consumed these and died and especially when no alternative food was available. In the laboratory trials, mice were caged individually and had easy access to alternative food as well as the ZnP-coated grains giving them ample opportunity to be selective in their choice of food. This contrasts with what occurs in a field situation where there is intraspecific competition for food and shelter and there is the threat of predation. The relationship between available alternative food and the efficacy of ZnP50 needs to be assessed experimentally in a field situation.

The higher values for ZnP LD_50_ provide us with a plausible explanation for the variable efficacy observed by farmers when the currently ZnP registered rate of 25 g ZnP/kg grain is applied. Our findings force us to conclude that ZnP bait applied at the currently prescribed rate of 1 kg/ha (25 g ZnP/kg grain) means that mice are not being presented with three lethal doses per square metre in the field (2-3 grains/m^2^). Instead, to be assured of receiving a lethal dose, mice would need to detect and consume at least two ZnP25-coated grains. This immediately raises questions about the capacity of mice to find a second or third ZnP25-coated grain prior to the onset of symptoms of toxicity in the complex environment in paddocks where conservation agriculture (CA) is practised.^22^ In our laboratory assessment where food was provided at the same time as the ZnP-coated grain, the proportion of mice dying was lower regardless of the ZnP dose per grain as mice ate fewer ZnP-coated grains (Table 3). The artificial nature of the laboratory comparison indicates that further comparative studies in CA cropping systems are required. We predict that the amount of alternative food available in the field at the time of ZnP baiting will influence the detection and uptake of ZnP-coated grains. In addition, the overall efficacy of a ZnP baiting program in field situations is likely to be affected by the growth stage of the crop being protected. In CA systems where available alternative food could be >200 grains/m^2^, this will likely affect ZnP baiting outcomes when the toxic grains are spread at only 2-3 grains/m^2^. Assessment of the efficacy of the higher dose (50 g ZnP/kg grain) compared to the registered dose at a population level under field conditions is needed to verify the findings of these laboratory-based trials.

## 5 CONCLUSIONS

This study has conclusively demonstrated that the LD_50_ for Australian wild house mice is more than two-fold higher than previously reported in the literature for wild mice. This indicates that the loading of ZnP on an individual grain should be doubled to achieve a lethal dose per grain and avoid bait aversion developing from sub-lethal effects of the toxin. Grains coated with the equivalent of an LD_90_ quantity (approximately 2 mg), were readily consumed by mice and mortality was greater than 90%, particularly in the absence of alternative food.

Results were less conclusive when alternative non-toxic grains were provided at the time of presentation of ZnP-coated grains. Assessment of the efficacy of the higher dose (50 g ZnP/kg grain) compared to the currently registered dose (25 g ZnP/kg grain) at a population level under field conditions is needed to validate the findings of these laboratory-based trials.

## ACKNOWLEDGEMENTS

This work was supported by the Grains Research and Development Corporation (GRDC) through project CSP1804-012RTX, and CSIRO Health and Biosecurity. We sincerely thank farmers for allowing us to trap mice on their properties for this experiment. We thank Leigh Nelson and Ken Young (GRDC) for ongoing support. Richard Duncan, University of Canberra, provided helpful advice on statistical analyses. All experiments and procedures were approved by the CSIRO Wildlife and Large Animal Ethics Committee (Approval No 2019-23) and conform to the Australian Code of Practice for the Care and Use of Animals for Scientific Purposes.

## DATA AVAILABILITY STATEMENT

The data that support the findings of this study are available from the corresponding author upon reasonable request.

---

**Table S1:** Time to death (h) after oral gavage with different doses of ZnP in oil for laboratory mice (Swiss Outbred), wild mice (Naïve) and wild mice (Exposed).

| Dose<br>(mg/kg) | n | No. humanely<br>killed | Time to death (h post gavage) |  |  |  |
| --- | --- | --- | --- | --- | --- | --- |
|  |  |  | Female<br>(fed) | Female<br>(unfed) | Male<br>(fed) | Male<br>(unfed) |
| **Outbred laboratory mice** |  |  |  |  |  |  |
| 50.22 | 8 | 2 | 25.0 |  | 25.3 |  |
| 65.1 | 6 | 2 | 31 |  | 48 |  |
| 74.4 | 5 | 0 |  |  |  |  |
| 83.7 | 8 | 5 | 9.25, 16.0 |  | 20.3, 29.5 | 17.5 |
| 102.3 | 8 | 7 | 8, 19.5 | 7.5 | 19, 23.5 | 10, 28 |
| 148.8 | 4 | 4 | 7 | 8.5 | 6.7 | 10 |
| **Wild mice (Naïve)** |  |  |  |  |  |  |
| 32.2 | 8 | 0 |  |  |  |  |
| 50.22 | 8 | 3 | 6.75 | 11.17 |  | 23.83 |
| 74.4 | 7 | 5 |  | 6.33 | 7.25, 9.5, 15.5 | 19 |
| 102.3 | 8 | 5 | 3.17, 3.42 |  | 3.25, 7.75 | 8.42 |
| 148.8 | 8 | 8 | 2.5 | 7, 22.33 | 4, 4.17, 8 | 8.17, 24.5 |
| **Wild mice (exposed)** |  |  |  |  |  |  |
| 30.41 | 8 | 0 |  |  |  |  |
| 50.22 | 7 | 1 |  |  |  | 24.0 |
| 65.1 | 4 | 1 | 19.83 |  |  |  |
| 74.4 | 8 | 5 | 5.67, 20.16 | 3.5 | 14.58 | 23.33 |
| 83.7 | 6 | 3 | 7.33, 32.67 | 6.67 |  |  |
| 102.3 | 11 | 10 | 5, 5.33, 6.75 | 4, 8.33 | 4.67, 5.25, 8 | 6, 24.0 |

**Table S2:** Summary of proportion of animals that died, modelled with dose, group, sex and fed or unfed as varying fixed effect factors. The top model included the oral gavage dose only (bold).

| Model | df | AIC | ΔAIC |
| --- | --- | --- | --- |
| **Dose** | **2** | **91.54** | 0 |
| Dose + Group | 6 | 97.23 | 5.69 |
| Dose + Group + Sex + Fed/Unfed | 8 | 97.97 | 6.43 |

**Table S3.** Number of mice consuming ZnP-coated grains and dying each day over 3 days, and overall mortality (%) for Trial 1 (T1) of each treatment. F25 = Fed, 25 g ZnP/kg grain; F50 = Fed, 50 g ZnP/kg grain; UF50 = UnFed, 50 g ZnP/kg grain. For F25 and F50 ZnP-coated grains and alternative standard wheat grains were added simultaneously. For UF50, alternative food was added 4 h later.

| Treatment | <u>Day 1</u> |  |  | <u>Day 2</u> |  |  | <u>Day 3</u> |  |  | <u>Mortality</u><br><u>over 3</u><br><u>days (%)</u> |
| --- | --- | --- | --- | --- | --- | --- | --- | --- | --- | --- |
|  | n | No.<br>taking | No.<br>dead<br>(%) | n | No.<br>taking | No.<br>dead | n | No.<br>taking | No.<br>dead |  |
| UF50 T1 | *9 | 9 | 8 (88.9) | 1 | 0 | 0 | 1 | 0 | 0 | 88.9 |
| F50 T1 | *9 | 9 | 5 (55.5) | 4 | 0 | 0 | 4 | 0 | 0 | 55.6 |
| F25 T1 | 10 | 10 | 5 (50) | 5 | 1 | 1 | 4 | 0 | 0 | 60.0 |

\*One animal in each group did not consume any ZnP-coated grains and was excluded from analysis.

**Table S4.**
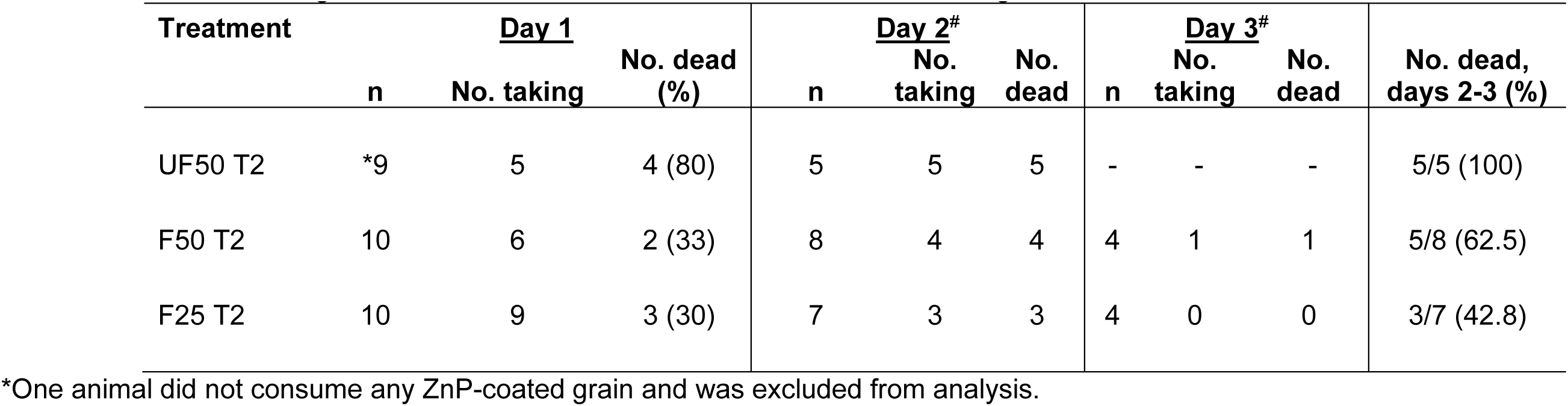
Number of mice consuming ZnP-coated grains and dying each day over 3 days, and overall mortality (%) for Trial 2 (T2) of each treatment. F25 = Fed, 25 g ZnP/kg grain; F50 = Fed, 50 g ZnP/kg grain; UF50 = UnFed, 50 g ZnP/kg grain. _#_ For Days 2 and 3, all groups were presented with ZnP-coated grains 6 h before addition of alternative standard wheat grains.

| Treatment | <u>Day 1</u> |  |  | <u>Day 2<sup>#</sup></u> |  |  | <u>Day 3<sup>#</sup></u> |  |  | No. dead,<br>days 2-3 (%) |
| --- | --- | --- | --- | --- | --- | --- | --- | --- | --- | --- |
|  | n | No. taking | No. dead<br>(%) | n | No.<br>taking | No.<br>dead | n | No.<br>taking | No.<br>dead |  |
| UF50 T2 | *9 | 5 | 4 (80) | 5 | 5 | 5 | - | - | - | 5/5 (100) |
| F50 T2 | 10 | 6 | 2 (33) | 8 | 4 | 4 | 4 | 1 | 1 | 5/8 (62.5) |
| F25 T2 | 10 | 9 | 3 (30) | 7 | 3 | 3 | 4 | 0 | 0 | 3/7 (42.8) |

\*One animal did not consume any ZnP-coated grain and was excluded from analysis.

